# Vaccinia virus attenuation by codon deoptimization of the A24R gene for vaccine development

**DOI:** 10.1101/2022.01.23.477337

**Authors:** María M Lorenzo, Aitor Nogales, Kevin Chiem, Rafael Blasco, Luis Martínez-Sobrido

## Abstract

Poxviruses have large DNA genomes and they are able to infect multiple vertebrate and invertebrate animals, including humans. Despite the eradication of smallpox, poxvirus infections still remain a significant public health concern. Vaccinia virus (VV) is the prototypic member in the *poxviridae* family and it has been used extensively for different therapeutic applications, including the generation of vaccines against multiple infectious diseases and/or for oncolytic treatment. Many attempts have been pursued to develop novel attenuated forms of VV with improved safety profiles for their implementation as vaccines and/or vaccines vectors. We and others have previously demonstrated how RNA viruses encoding codon-deoptimized viral genes are attenuated, immunogenic and able to protect, upon a single administration, against challenge with parental viruses. In this study, we employed the same experimental approach based on the use of misrepresented codons for the generation of a recombinant (r)VV encoding a codon-deoptimized A24R gene, which is a key component of the viral RNA polymerase. Similar to our previous studies with RNA viruses, the A24R codon-deoptimized rVV (v-A24cd) was highly attenuated *in vivo* but able to protect, after a single intranasal dose administration, against an otherwise lethal challenge with parental VV. These results indicate that poxviruses can be effectively attenuated by synonymous codon deoptimization and open the possibility of using this methodology alone or in combination with other experimental approaches for the development of attenuated vaccines for the treatment of poxvirus infection, or to generate improved VV-based vectors. Moreover, this approach could be applied to other DNA viruses.

**IMPORTANCE:** The family *poxviridae* includes multiple viruses of medical and veterinary relevance, being vaccinia virus (VV) the prototypic member in the family. VV was used during the smallpox vaccination campaign to eradicate variola virus (VARV), which is considered a credible bioterrorism threat. Because of novel innovations in genetic engineering and vaccine technology, VV has gained popularity as a viral vector for the development of vaccines against several infectious diseases. Several approaches have been used to generate attenuated VV for its implementation as vaccine and/or vaccine vector. Here, we generated a rVV containing a codon-deoptimized A24R gene (v-A24cd), which encodes a key component of the viral RNA polymerase. v-A24cd was stable in culture cells and highly attenuated *in vivo* but able to protect against a subsequent lethal challenge with parental VV. Our findings support the use of this approach for the development of safe, stable, and protective live-attenuated VV and/or vaccine vectors.

## INTRODUCTION

Poxviruses belong to a family of large, double stranded DNA viruses, designated *Poxviridae*, which replicate and assemble entirely in the cytoplasm of infected cells (1–3). Poxviruses are able to infect a broad range of both invertebrate and vertebrate animals, including humans and wildlife or domestic animals, and cause disease in many of them (4–8). Therefore, they are considered an important threat to public human health (6–8).

Smallpox is caused by variola virus (VARV) which, together with vaccinia virus (VV), belong to the orthopoxvirus genus, and constituted one of the deadliest diseases in human history that killed approximately 300 million people only in the twentieth century (9–11). Thanks to a world-wide vaccination program, the lack of non-human reservoirs, and virus slow mutation rate, the World Health Organization (WHO) declared smallpox eradicated in 1980 (11, 12). However, worries remains because the possibility that smallpox may re-emerge accidently from forgotten stocks of VARV or even from *de novo* synthesis by using current biotechnologies (10, 13). In fact, the United States (US) military considers the risk of weaponized smallpox sufficient to justify the continued vaccination of all its military personal against the disease (14). Given the low worldwide vaccination rate against smallpox since 1980, population could be highly vulnerable to a new outbreak of smallpox, increasing the current concerns (10, 13, 15). Furthermore, with increasing global trade many poxviruses infecting animals could have an important impact on commercially relevant livestock and ecologically endangered wildlife (4–8, 14). A particularly relevant case is monkeypox which causes a disease with symptoms similar to, but less severe than, smallpox. Monkeypox is a zoonosis with different hosts and human-to-human transmission is limited. However, fifteen countries on four continents have reported confirmed human cases of monkeypox. Most are usually found near tropical rainforests, where there are animals carrying the virus, but there have been emergencies in other remote countries such as the US and the United Kingdom. Importantly, there is currently no specific treatment for monkeypox (16), although vaccination against smallpox has been shown to be approximately 85% effective in preventing monkeypox (17, 18).

VV was used to prevent and eradicate smallpox and it has become the prototypic member of the *poxviridae* family. Importantly, novel genetic engineering methods to facilitate the manipulation of the genome of poxvirus (19–22) have allowed researchers to use VV as a vaccine vector for the development of vaccines against a variety of infectious diseases, and/or for oncolytic treatment (9, 23–28). In addition, poxviruses, mainly VV, have also multiple biotechnological applications based on heterologous gene expression (19).

A key factor to the optimization of poxviruses as vaccine vectors is the balance between the safety profile and immunogenicity *versus* retained virulence. Although the smallpox vaccine is generally considered safe and VV is naturally attenuated, classical VV vaccine strains still may produce some complications in patients with systemic immune deficits, which should be exempt from current vaccination programs because of significant increased risks of adverse or severe outcomes (15, 29, 30). Therefore, several strategies have been applied to increase the attenuation of poxviruses as vaccine vectors (9, 23, 26). In general those include deletion of genes and point mutations as a result of extensive passaging in cell culture (e.g. modified vaccinia Ankara, MVA) (22, 31–34) or the introduction of multiple defined deletions (NYVAC) (9, 35). Alternatively avipoxviruses, like fowlpox and canarypox, which are not well adapted to mammalian cells, have been used as vaccine vectors since they are not able to undergo full replication in mammalian cells (7, 9, 26). Therefore, although a variety of attenuated poxviral vaccine vectors are currently available they have been derived from blind processes.

The mammalian genetic code is degenerated, and most amino acids are coded by multiple synonymous codons, which are not used with equal frequency within or between DNA/RNA genomes (36, 37), showing differences in the frequency at which organisms use codons to incorporate the same amino acid residue into a protein (38–40). This biological phenomenon called codon usage bias has been used for codon optimization (CO) or codon deoptimization (CD) approaches to increase or decrease, respectively, gene expression in different expression systems. While CO, where each amino acid is encoded by the most abundant codon, is widely employed to increase protein expression for pharmaceutical, biotechnological or research purposes (41–45), the potential of CD has been less explored (46–52). CD is achieved by replacing original codons for those with less-preferred usage (46–50). Importantly, changes are only at the nucleotide level without affecting the amino acid sequence of the protein or their immunogenic properties and functionality (53, 54). We and others have previously documented the feasibility of generating recombinant RNA viruses, harboring codon deoptimized genes, with attenuated phenotypes as potential vaccines and/or vaccine vectors (46–52). For that, several genes of RNA viruses as influenza A virus (IAV) (46, 47), the arenaviruses lymphocytic choriomeningitis virus (LCMV) or Lassa virus (LASV) (48–50), and respiratory syncytial virus (RSV) (52) were CD, and the *in vitro* and *in vivo* characteristics of the recovered viruses were analyzed in different *in vitro* and *in vivo* animal models. However, to date, the use of a CD-based approach for the development of attenuated forms of viruses for their implementation as safe and protective live-attenuated vaccines have not been explored for DNA viruses.

Since VV encodes all the enzymes required for DNA replication and transcription, virus replication occurs independently of the host. One key virally-encoded enzyme is the viral RNA polymerase, which is required throughout the replication cycle for transcription of viral genes. In this work, we have reengineered the VV A24R gene, encoding the second largest subunit of the viral RNA polymerase rpo132 (55–59), by introducing through *de novo* gene synthesis, the least used human synonymous codons without modifying the viral protein amino acid sequence. The generated recombinant virus containing synonymous CD mutations in the A24R gene (v-A24cd) showed lower replication levels and altered plaque phenotype in culture cells. Moreover, v-A24cd was highly attenuated in a mouse model of infection. Importantly, immunization of mice using a single intranasal dose of v-A24cd conferred full protection against a lethal challenge with parental VV, demonstrating the feasibility of using this CD-based approach for the development of novel and safer VV vaccines and/or vaccine vectors, or for the its implementation to attenuate other DNA viruses.

## RESULTS

### CD of VV A24R gene results in reduced levels of protein expression

Codon deoptimization (CD) represents a potential strategy to modify the expression efficiency of a gene, thereby altering the amount of its protein product. We selected the A24R gene as a target to examine the potential of using our CD-based approach with VV for the generation of a live-attenuated vaccine and to demonstrate the feasibly of implementing this CD-based strategy for the attenuation of DNA viruses. A24R encodes the catalytic subunit of VV RNA polymerase, which plays a critical role in viral gene transcription (55–59).

Therefore, the CD of VV A24R gene, most likely, will affect the expression levels of other viral proteins, disrupting multiple steps during VV infection. To evaluate the effect of CD on VV A24R expression, we engineered its open reading frame (ORF) using underrepresented codons in human cells but preserving the intact A24R amino acid sequence. The CD A24R (A24Rcd) sequence (**Figure 1A** **and supplementary** **Figure 1**) included 666 (57.17%) codon changes through 775 (22.17%) nucleotide substitutions (**Figure 1B**) and the gene was synthesized and cloned in plasmids suited for insertion into the VV genome (**Figure 2**). In our design, we included the A24R versions and recombination flanks for the A24R gene L and R (left and right) for the generation of rVV by homologous recombination. In addition, a GFP expression cassette was included for easy isolation and visualization of recombinant viruses (**Figure 2**).

**Figure 1.**
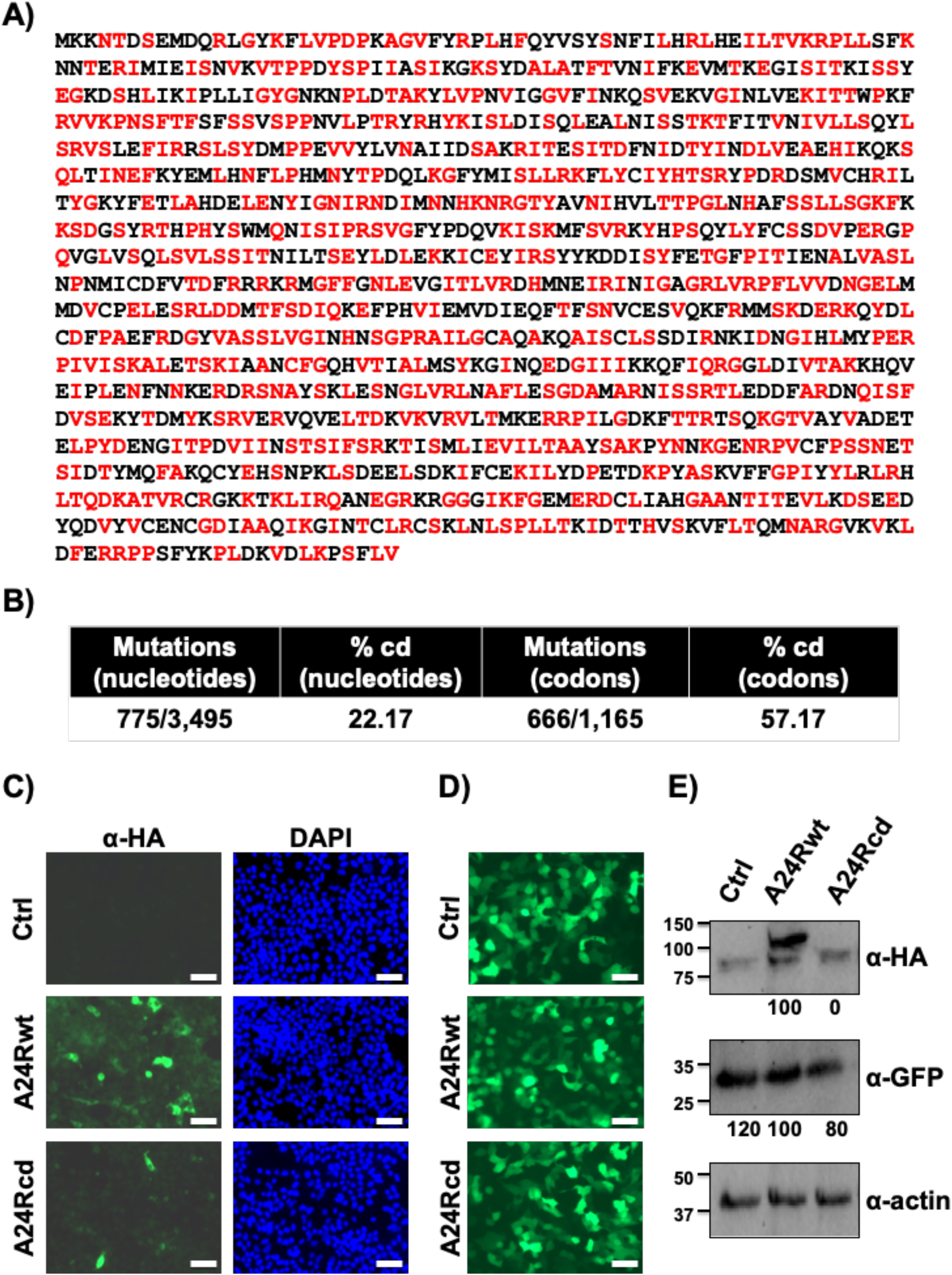
Codon deoptimization reduces VV A24R expression levels. **A**) **Amino acid sequence of A24R.** Codon-deoptimized amino acid residues are indicated in red (see **Supplementary** Figure 1 for wild-type and codon-deoptimized nucleotide sequences). Methionine (M) and tryptophan (W) amino acid residues, as well as amino acids already associated with deoptimized codons are indicated in black. **B**) **Mutations in A24Rcd.** Number and percentage of nucleotides or codons changes in the codon-deoptimized A24R are indicated. **C**) **Analysis of protein expression in transfected cells by immunofluorescence.** Human 293T cells were transfected with 1 μg of pCAGGS expression plasmids encoding C-terminal HA-tagged A24Rwt or A24Rcd proteins. At 48 h p.t., protein expression was assessed by immunofluorescence using an anti-HA pAb. DAPI was used for nuclear staining. Scale bars, 100 µm. **D-E**) **Analysis of protein expression by Western blot.** Human 293T cells were transiently co-transfected with pCAGGS expression plasmids encoding C-terminal HA-tagged versions of A24Rwt or A24Rcd proteins (2 μg) together with a pCAGGS plasmid encoding GFP (0.5 μg). At 48 h p.t., GFP expression was assessed under a fluorescence microscope (**D**). A24Rwt or A24Rcd protein expression levels were assessed by Western blot using an anti-HA pAb (**E**). GFP expression levels were assessed using an anti-GFP pAb. Actin expression levels were determined using an anti-actin MAb and were used as loading controls. Numbers in the left indicate the size of molecular markers in kDa. Scale bars, 200 µm. Western blots were quantified by densitometry using the software ImageJ. Relative band intensities as described in Materials and Methods are indicated.

**Figure 2.**
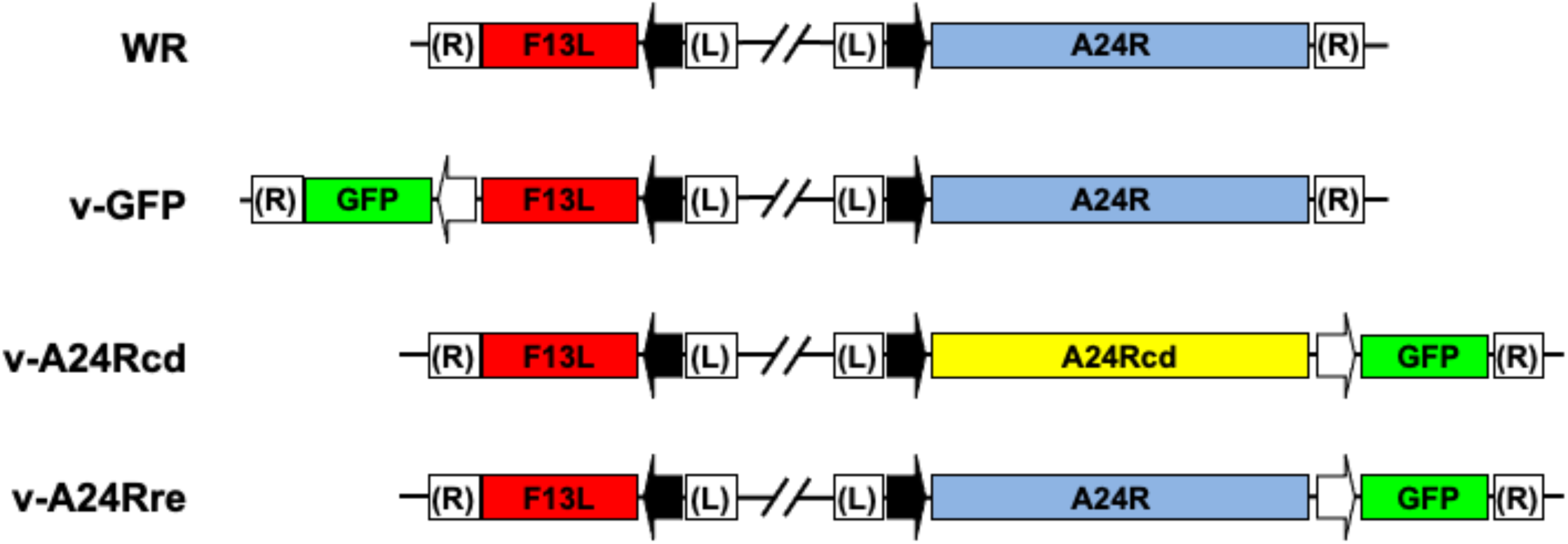
Schematic representation of rVV. The names and the viral genomic organization around the F13L and A24R loci of the rVV used in this study are indicated. Viral promoters (black arrows for F13 and A24R and white arrows for GFP), as well as genes encoding for GFP (green boxes), F13L (red boxes) and WR, and codon deoptimized (cd) and revertant (re) A24R (yellow and blue boxes, respectively) are shown.

To determine whether CD of A24R could lead to a reduction in protein expression levels in the absence of infection, HA-tagged A24Rwt or A24Rcd ORF sequences were cloned into pCAGGS expression plasmids. Then, human 293T cells were transiently transfected with pCAGGS-A24Rwt-HA or pCAGGS-A24Rcd-HA constructs, and protein expression was analyzed by immunofluorescence using an antibody against the HA epitope tag (**Figure 1C**). A significant reduction in both the fluorescent signal and the number of fluorescent cells was observed in cells transfected with the plasmid encoding A24Rcd-HA as compared to cells transfected with the A24Rwt-HA plasmid (**Figure 1C**). The effects of CD on A24R protein expression were also assessed by Western blot. For that, human 293T cells were transiently co-transfected with pCAGGS plasmids expressing GFP and A24Rwt or A24Rcd genes and protein expression was evaluated at 48 hours post-transfection (h p.t.) GFP examination using a fluorescence microscope (**Figure 1D**) or by Western blot (**Figure 1E**) confirmed that A24R (A24Rwt or A24Rcd) did not affect GFP expression, indicating that A24R sequence is not toxic in transfected cells, and the observed reduced expression of A24Rcd is due to the sequence of codons used. Whereas A24R was clearly detected, the level of expression of A24Rcd was drastically reduced, in agreement with the fluorescence results (**Figure 1E**). Protein densitometry of Western blot bands further confirmed this observation. These data indicate that, similar to the proteins of RNA viruses, codon deoptimization of VV A24R reduces protein expression in mammalian cells.

### Generation and growth properties of v-A24Rcd in cell culture

To evaluate the effect of A24R CD in viral fitness and pathogenicity, we next used well-established poxvirus genetics techniques to isolate viruses expressing modified versions of A24R (**Figure 2**). Substitution of the A24R gene by a CD version was carried out by insertion of an A24 CD/GFP construct in place of the normal A24R gene, to obtain virus v-A24cd. A revertant virus expressing the normal A24R gene (v-A24Rre) was subsequently obtained from vA24cd by recombination of an A24/GFP construct into the same locus. As an additional control, we used vGFP, which contains a GFP cassette at a distant place in the genome and has an unmodified A24R gene (**Figure 2**).

Next, we assessed the replication properties of the generated rVV by multicycle growth kinetics in BSC-1 cells infected at low multiplicity of infection (MOI, 0.01). Progress of infection was followed by examining GFP expression under a fluorescence microscope (**Figure 3A**) or the presence of infectivity in cell culture supernatants using plaque assays (**Figure 3B**). The recombinant v-GFP and v-A24Rre displayed similar levels of GFP expression and replication kinetics (**Figures 3A and 3B**, respectively) and significant differences were not observed between them. On the other hand, GFP expression from v-A24Rcd was considerably reduced (**Figure 3A**) and the virus did not achieve replication levels similar to those of v-GFP and v-A24Rre (**Figure 3B**). In addition, we measured the presence of intracellular and extracellular v-GFP, v-A24Rcd and v-A24Rre in BSC-1 infected cells (MOI, 3) at 24 hours post-infection, h p.i. (**Figure 3C**). We observed a decrease in cell-associated and extracellular v-A24Rcd titers as compared with v-GFP and v-A24Rre, suggesting that our CD-based strategy has a general effect in the production of progeny virus. We also examined the phenotype of v-A24Rcd, v-A24Rre or v-GFP in BSC-1 cells using plaque assays and GFP expression (**Figure 3D**). As expected, based on our multicycle growth kinetics (**Figures 3A and 3B**), v-A24cd produced significantly smaller plaques than the control recombinant viruses vGFP and v-A24Rre (**Figure 3D**). Altogether, these data demonstrates that growth of v-A24Rcd was impaired as compared to v-GFP or v-A24Rre in cultured cells.

**Figure 3.**
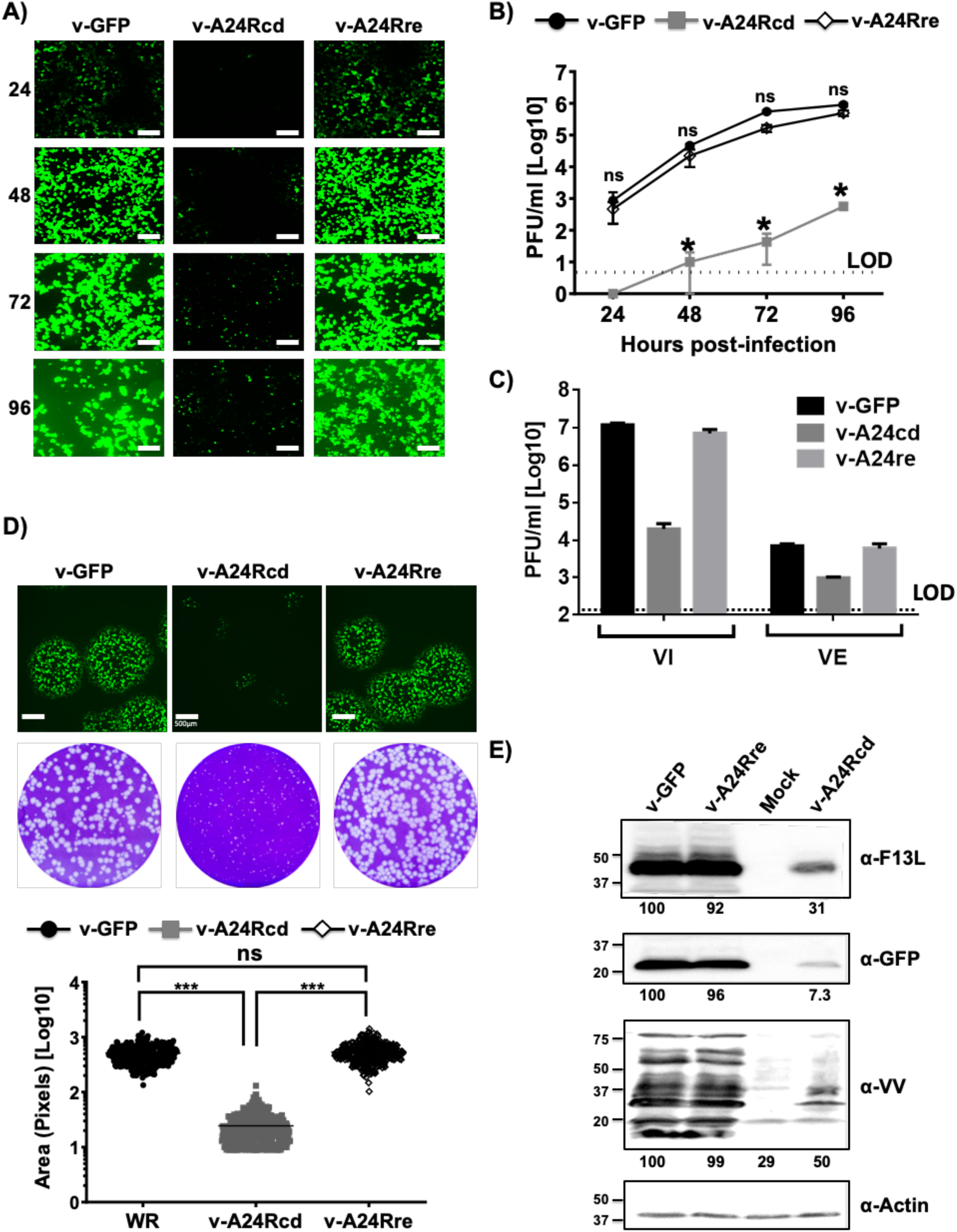
Characterization of v-A24Rcd. **A-B**) **Growth kinetics.** BSC-1 cells monolayers (6-well plate format, 10^6^ cells/well) were infected (MOI 0.01) in triplicate with the indicated viruses and at the indicated h p.i., GFP fluorescence was detected by fluorescence microscopy (**A**). Scale bars, 200 µm. Cell culture supernatants were also collected at the indicated times p.i. and viral titers were determined using standard plaque assay (**B**). *, P < 0.05, using Student’s t test (v-GFP versus v-A24Rcd; not significant (ns) differences were obtained between v-GFP and v-A24Rre). **C**) **Production of extracellular and cell-associated virus.** BSC-1 cells (6-well plate format, 10^6^ cells/well) were infected at an MOI of 3 PFU in triplicate. At 36 h p.i., virus in the cell culture medium (VE) and associated with the cells (VI) was titrated by plaque assay on fresh BSC-1 monolayers. **D**) **Plaque phenotype.** BSC-1 cell monolayers (6-well plate format, 10^6^ cells/well, triplicates) were infected with ∼100-200 PFU/well of the indicated viruses and incubated with medium containing methyl cellulose. At 3 days p.i., monolayers were fixed, GFP fluorescence was imaged (top) and wells were stained with crystal violet (middle). Scale bars, 100 µm. The area of plaques of 200 viruses was quantified and is represented (bottom). ***, P < 0.0001, using Student’s t test. ns: not significant differences. **E**) **Protein expression levels.** BSC-1 cells (6-well plate format, 10^6^ cells/well) were either mock-infected or infected (MOI 3) with the indicated viruses and viral protein expression levels were analyzed at 24 h p.i. by Western blot with antibodies against F13L, GFP or a pAb against VV. Actin was used as loading control. Western blots were quantified by densitometry using the software ImageJ. Bands were normalized to actin expression. Numbers in the left indicate the size of molecular markers in kDa.

To evaluate if infection with v-A24Rcd led to differential expression of viral proteins, lysates of infected BSC-1 cells were evaluated by Western blot (**Figure 3E**). To that end, cells were either mock-infected or infected (MOI, 3) and cell extracts were collected at 24 h p.i. and probed using specific antibodies against VV F13 protein, GFP, and VV. An antibody against actin was used as a protein loading control. The levels of F13, GFP, and viral proteins were not altered significantly between v-GFP or v-A24Rre infected cells. However, F13, GFP and total VV protein expression levels were notably reduced, in v-A24Rcd-infected BSC-1 cells as compared to v-GFP or v-A24Rre. Altogether, these data suggest that v-A24Rcd results in an expected decrease in viral protein expression levels in infected cells, that appears as a general effect on viral protein accumulation as revealed with anti-VV polyclonal antiserum (**Figure 3E**).

### Viral factory formation and subcellular localization of B5 protein

Previous experiments show a block in virus replication as well as a decrease in viral protein accumulated during infection with v-A24cd (**Figure 3**). To characterize in more detail the infection by v-A24cd, we studied the last steps in the virus life cycle by fluorescence microscopy (**Figure 4**). The results confirmed a decrease in GFP expression, and a reduction in the number of viral factories in vA24cd-infected cells when compared to those infected with the control viruses v-GFP and vA24re (**Figure 4**). In addition, we visualized by immunofluorescence the distribution of protein B5, which is localized in the Golgi complex, where it is incorporated into infectious virus that are then transported and released into the extracellular medium (60). B5 labeling was decreased in v-A24cd infected cells, being barely detectable at 8 h p.i. (**Figure 4A**). However, although at later times (18 h p.i., **Figure 4B**) the amount of B5 was also low, its distribution appeared normal, being accumulated in a juxtanuclear region consistent with the Golgi complex and puncta representing enveloped virus particles. These results are indicative of a virus replication cycle that is delayed with respect to normal virus, likely as a result of low protein expression resulting from the CD of A24R.

**Figure 4.**
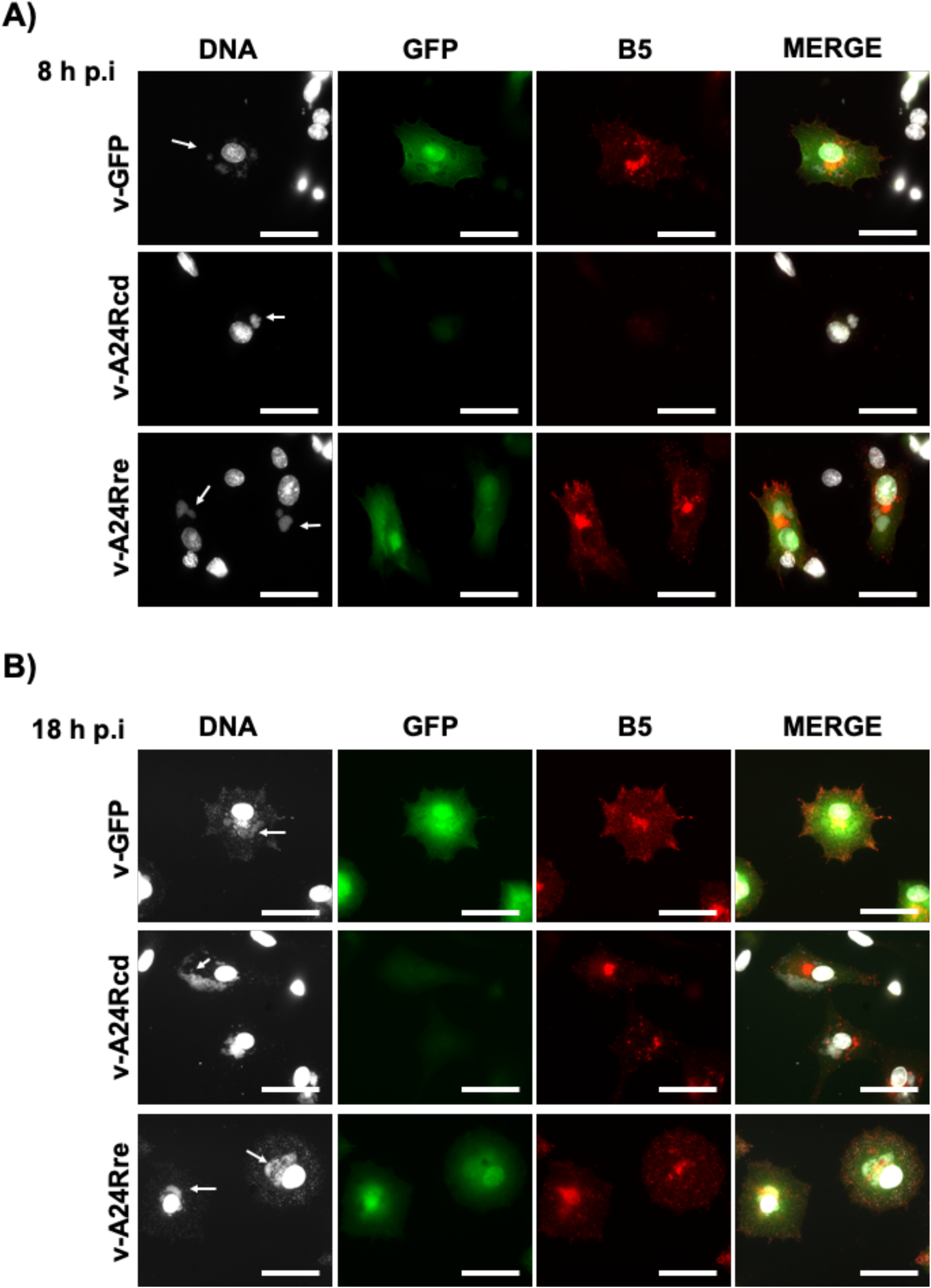
Subcellular localization of B5 in infected cells. Monolayers of BHK-21 cells grown on cell coverslips were infected with the virus indicated on the left. At 8 h p.i. (**A**) or 18 h p.i. (**B**) cells were fixed, and subjected to immunofluorescence with anti-B5 antibody. For visualization of viral factories (arrows), cells were incubated with Hoechst staining. Scale bars, 50 µm.

### *In vivo* characterization of v-A24Rcd

Because we observed that replication properties and viral protein expression of v-A24Rcd were affected in BSC-1 cells (**Figure 3**), including viral factory formation (**Figure 4**), we postulated that v-A24Rcd would be attenuated *in vivo*. To demonstrate this hypothesis, we evaluated and compared the virulence of v-A24Rcd, v-A24Rre and parental viruses in a mouse model of viral infection (**Figure 5**). For that, groups of C57BL/6 mice (n = 6/group) were inoculated intranasally (i.n.) with 10^4^, 10^5^, or 10^6^ plaque forming units (PFU)/animal of v-A24Rcd, v-A24Rre or WR and mice were monitored during 14 days for body weight loss (**Figure 5A**) and mortality (**Figure 5B**). In addition, a control group was mock-infected (PBS). Mice infected with v-A24Rre or parental WR viruses showed clear and similar changes in morbidity (**Figure 5A**) and all succumbed to viral infection independently of the viral dose (**Figure 5B**). Notably, v-A24Rcd was highly attenuated, with no changes in body weight and all the animals surviving viral infection (**Figure 5B**), even with the highest dose of 10^6^ PFU.

**Figure 5.**
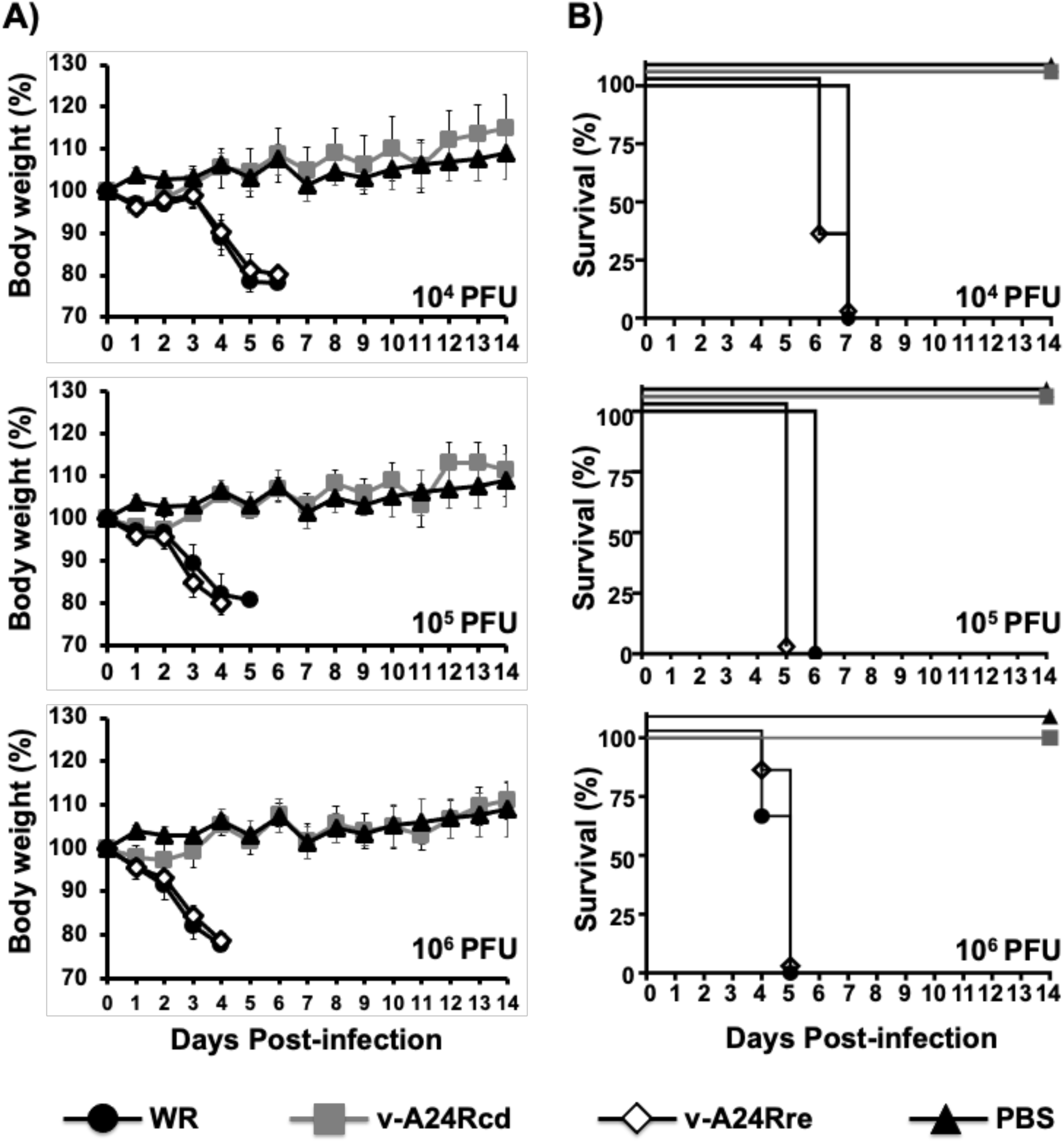
*In vivo* attenuation of v-A24Rcd. Six-to-eight-week-old female C57BL/6 mice (n = 6/group) were mock (PBS)-infected or infected intranasally with 10^4^, 10^5^, or 10^6^ PFU of VV WR, v-A24Rcd or v-A24Rre. Morbidity (**A**) and survival (**B**) were evaluated for 14 days. Mice that lost 25% of their initial body weight were sacrificed. Data represent the means and SD of the results determined for individual mice.

### v-A24Rcd protects mice from lethal viral challenge with parental VV

Despite the fact that v-A24Rcd was highly attenuated in mice (**Figure 5**), we hypothesized that v-A24Rcd could protect against another wise subsequent lethal challenge with parental VV. To assess this, animals inoculated with 10^4^, 10^5^, or 10^6^ PFUs/animal of v-A24Rcd, or mock-vaccinated, (**Figure 5**) were challenged 21 days later with 10^4^ PFU/mice of parental VV WR and morbidity (**Figure 6A**) and mortality (**Figure 6B**) were monitored for 2 weeks. None of the v-A24Rcd-vaccinated animals, independently of the vaccination dose, showed any changes in body weight loss (**Figure 6A**), and all animals survived the lethal challenge with parental VV (**Figure 6B**). Contrary, all mock-vaccinated mice drastically lost weight and died after parental VV WR challenge (**Figures 6A and 6B**, respectively). These results indicate that a single dose of v-A24Rcd is able to protect against a lethal challenge with parental VV WR, demonstrating the feasibility of its use as a safe and protective live-attenuated vaccine, or vaccine vector.

**Figure 6.**
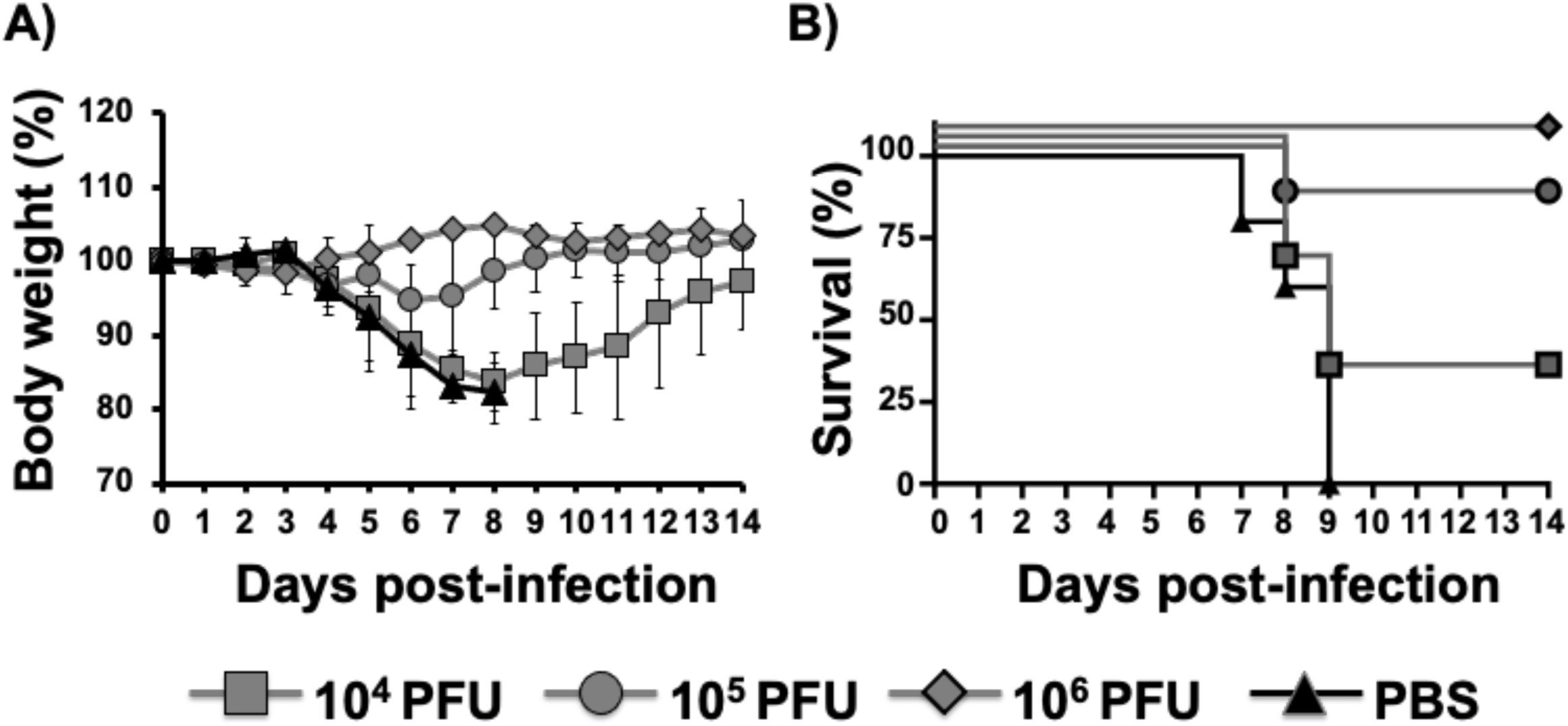
Protection efficacy of v-A24Rcd. Six-to-eight-week-old female C57BL/6 mice (n = 6/group) were mock (PBS)-vaccinated or vaccinated with 10^4^, 10^5^, or 10^6^ PFU of v-A24Rcd (mice from Figure 5). Twenty-two days after vaccination, mice were challenged intranasally with 10^4^ PFU/mice of VV WR and morbidity (**A**) and survival (**B**) were evaluated for 2 weeks. Data represent the means and SD of the results determined for individual mice.

### Stability of v-A24Rcd in culture cells

A critical concern with live-attenuated vaccines or vaccine vectors is the prospect of genetic instability that might lead to the loss of attenuation. To assess stability of v-A24Rcd, we serially passaged the virus in BSC-1 cells for a total of 10 passages (**Figure 7**). Next, virus growth properties from the first (v-A24Rcd.P1) and last (v-A24Rcd.P10) passage were evaluated by plaque assay (**Figures 7A-7C**) and multicycle growth kinetics (**Figure 7D**). For these assays, v-GFP and parental viruses were also included as controls. Notably, results indicate that v-A24Rcd was phenotypically stable since both v-A24Rcd.P1 and v-A24Rcd.P10 displayed the same plaque phenotype (**Figures 7A and 7B**) and levels of GFP expression (**Figure 7A**), with A24Rcd.P1 and v-A24Rcd.P10 producing smaller plaques than v-GFP or parental viruses (**Figure 7C**). In addition, v-A24Rcd.P1 and v-A24Rcd.P10 grew with indistinguishable kinetics reaching the same viral titers that were both significantly reduced as compared to v-GFP or parental viruses (**Figure 7D**). Altogether, these results suggest that v-A24Rcd is phenotypically stable *in vitro*, which is an important feature for the implementation of v-A24Rcd as a live-attenuated vaccine and/or vaccine vector.

**Figure 7.**
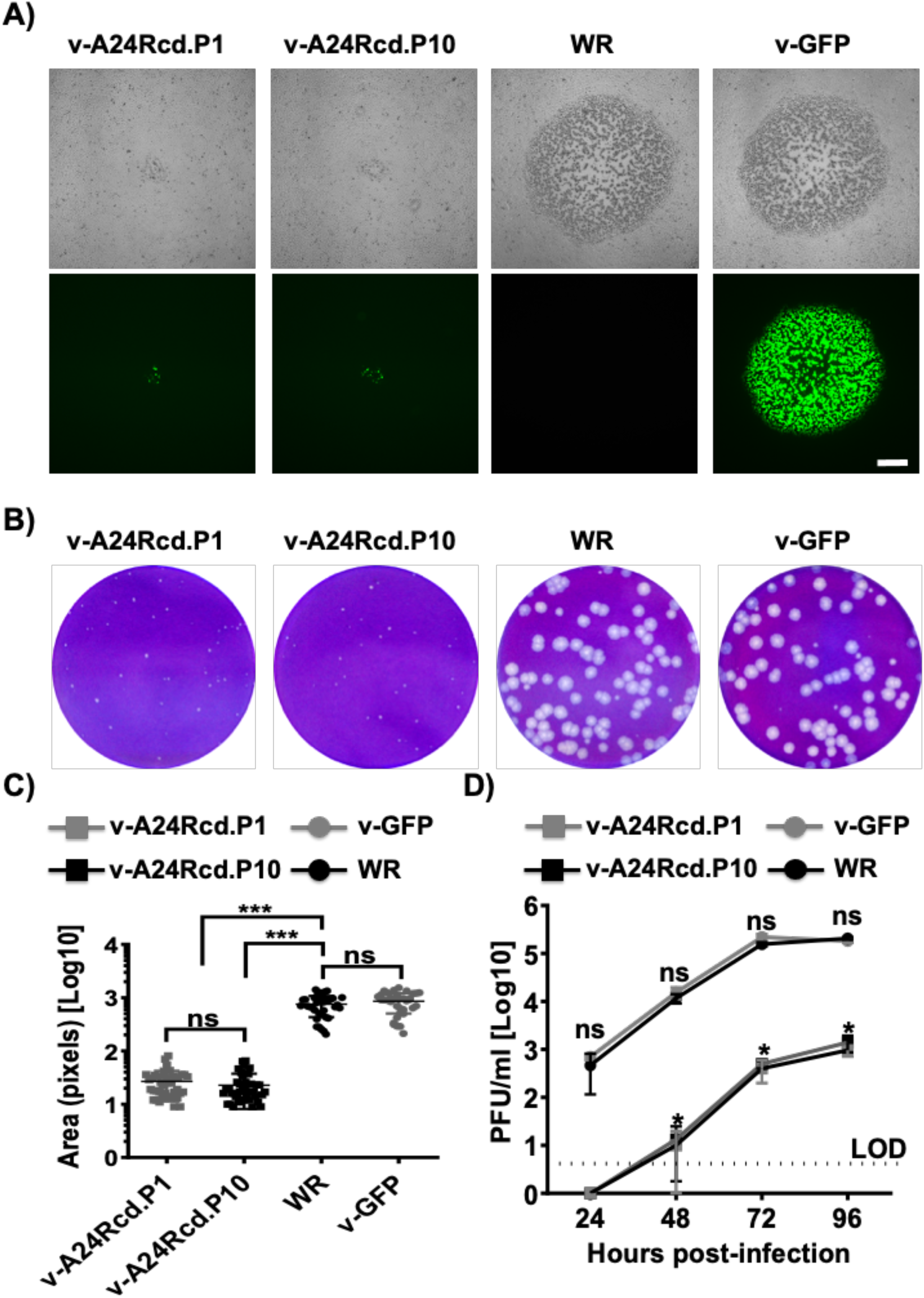
Stability of v-A24Rcd. v-A24Rcd was subjected to 10 serial passages in BSC-1 cell monolayers (6-well plate format, 10^6^ cells/well). Virus from the first (v-A24Rcd.P1) and last (v-A24Rcd.P10) passages was characterized by plaque assay and viral growth kinetics. **A**) **Plaque phenotype.** Viral plaques of v-A24Rcd.P1, v-A24Rcd.P10, WR and v-GFP infected BSC-1 cells (6-well plate format, 10^6^ cells/well) were visualized by bright microcopy (top) and GFP expression (bottom). Scale bars, 500 µm. **B-C**) **Plaque size.** Monolayers of BSC-1 cells (6-well plate format, 10^6^ cells/well) were inoculated with 50-100 PFU of the indicated viruses and incubated with medium containing methyl cellulose. Virus plaques 3 days after infection were visualized by crystal violet staining (**B**) and the area of plaques (n=30) was measured (**C**) ***, P < 0.0001, using Student’s t test. ns: not significant differences. **D**) **Growth kinetics.** BSC-1 cells (6-well plate format, 10^6^ cells/well) were infected (MOI 0.01) in triplicates with the indicated viruses and viral titers in cell culture supernatants from infected cells at the indicated times p.i. were determined by plaque assay. *, P < 0.05, using Student’s t test (WR versus v-A24Rcd.P1 or P10; no significant differences were obtained between WR and v-GFP, or v-A24Rcd.P1 and P10). ns: not significant differences.

## DISCUSSION

The accumulated knowledge of poxviruses, including the prototype VV, together with the development of recombinant DNA techniques, have expanded the use of poxviruses as vaccine vectors to prevent infections in humans and domestic or wild animals (7, 9, 19, 23, 24, 26, 27). VV offers many advantages as vaccine vector, including low virulence, induction of good immune responses, and large insertion capacity and stability. However, the virus retains some degree of virulence and may cause clinical complications in immunocompromised individuals (15, 29, 30). Thus, strategies to attenuate the virus are sought. Multiple approaches have been developed to date to increase the attenuation of poxviruses as vaccine vectors including the deletion of viral genes or genome fragments affecting different steps in viral infection (9, 23, 26). In this report, we describe, for the first time, the feasibility of using a CD-based approach for the development of attenuated forms of VV that can be used as live-attenuated vaccines for the treatment of poxvirus infections, or vaccine vectors.

Protein expression of mammalian viruses is influenced by the codon usage bias of the cells they infect, and the use of suboptimal codons has gained attention recently as a novel strategy for the development of viral live-attenuated vaccines (46–50). It has been shown that the CD of genes from RNA viruses leads to a reduction of viral protein expression, which usually is associated to defects in viral replication and, consequently, *in vivo* attenuation (46–52). Although, protein translation efficiency seems to be a major factor, we cannot rule out that other factors may be responsible for viral attenuation, including changes in RNA secondary structure, stability, and/or gene composition (e.g.: %GC content) (36, 40, 53, 61). In any event, CD of viral genes in a number of systems have resulted in attenuated forms of the viruses with an excellent safety and protective profile.

In this work, a rVV was designed under the hypothesis that introduction of many underrepresented codons (CD) into the key A24R gene should lead to a reduction of viral replication and protein expression levels and thus an attenuated phenotype. We selected A24R for our initial studies to explore the effect of CD because A24R gene encodes one of the constituents of the viral RNA polymerase core complex and therefore it is involved in the transcription of other viral genes (55–59). Thus, the decrease in the amount of RNA polymerase can lead to a reduction in the synthesis of other viral proteins. According to our results, CD of A24R resulted in reduced levels of protein expression, as compared to A24R WT, in transfected cells (**Figure 1**). Notably, a rVV harboring an entire A24Rcd gene, v-A24Rcd (**Figure 2**), showed reduced replication levels in culture cells (**Figure 3**) that were also associated to reduced expression of viral proteins (**Figure 3**), including a reporter gene (GFP) that was inserted in the viral genome to easily track viral replication. The easiest explanation is that reduced A24 protein levels lead to a general reduction of late protein production, thereby affecting viral progeny levels (**Figure 4**).

The *in vitro* results readily translated to *in vivo*, where v-A24Rcd was highly attenuated compared to the parental (WR) or revertant (v-A24Rre) viruses (**Figure 5**). Importantly, we found that although v-A24Rcd was highly attenuated in mice, vaccination with a single dose of v-A24Rcd protected mice against a lethal challenge with parental VV (**Figure 6**). Moreover, our viral stability studies indicate that v-A24Rcd was phenotypically stable *in vitro* (**Figure 7**). These results provide proof of concept for using a CD-based approach as a valid strategy to modulate poxvirus virulence and the development of poxvirus-based vaccines and/or vaccine vectors. In fact, recombinant virus containing several CD genes involved in different steps of the virus life cycle could be used to generate novel vaccines and/or vaccines vectors with different levels of attenuation. Moreover, our studies provide the first evidence that a CD approach can be applied to large DNA viruses, as we previously described with RNA viruses (46–50), and that viral attenuation induced by CD is most likely associated, at least in part, to reduced levels of protein expression during viral infection. Moreover, and given that the A24R gene we inserted in v-A24Rcd was fully deoptimized, this strategy may lead to a way to custom-define the degree of deoptimization by choosing an intermediate level of codon deoptimization, as previously described for RNA viruses (48).

Notably, a previous study by Eschke *et al*. (62) using a codon pair bias deoptimization (CPBD) approach which has been also efficient for attenuation of RNA viruses (47, 63–68), suggest the feasibility of altering the codon usage of large DNA viruses, e.g. Marek’s disease virus (MDV). CPBD relies on recoding of genes by increasing the number of codon pairs that are underrepresented in the host without changing the amino acid sequence of the encoded proteins. Therefore, both CPBD and CD strategies could decrease protein production of functional genes, affecting viral fitness and causing viral attenuation. Authors, recoded UL30 that encodes the catalytic subunit of the viral DNA polymerase to minimize the level of overrepresented codon pairs of the MDV host, the chicken. A fully CPBD UL30 mutant virus was not recovered, although authors were able to recover recombinant viruses containing partially CPBD UL23 genes. In cultured cells, the partially CPBD UL23 MDV shown reduced levels of protein expression and lower titers compared to parental virus. Moreover, partially CPBD UL23 MDV were attenuated *in vivo*, indicating that CPBD could also be used as a suitable approach to attenuate large DNA viruses by recoding their genome. These results, together with those in this manuscript, demonstrate the feasibility of using CPBD- or CD-based approaches to generate attenuated forms of DNA viruses (MDV and VV, respectively), as potential live-attenuated vaccines for the treatment of DNA virus infections and/or attenuated vaccine vectors for the treatment of other human diseases.

## MATERIALS AND METHODS

### Cells and viruses

Human embryonic kidney (HEK) 293T (American Type Culture Collection, ATCC, CRL-11268) and African green monkey kidney epithelial BSC-1 (ATCC CCL-26) cells were cultured and maintained in Dulbecco’s modified Eagle’s medium (DMEM, 293Tcells) or in Eagle’s minimal essential medium (EMEM, BSC-1 cells) supplemented with 5% fetal bovine serum (FBS) and 1% PSG (penicillin, 100 units/mL; streptomycin 100 μg/mL; L-glutamine, 2 mM) at 37 °C with 5% CO2.

Vaccinia virus (VV) Western Reserve (WR) strain was obtained from the American Type Culture Collection (ATCC VR-119). The rVV expressing the green fluorescent protein (v-GFP) was previously described (20–22).

### Construction of plasmids containing codon-deoptimized A24R

To engineer a rVV containing the A24R codon-deoptimized gene (v-A24Rcd) through genome-scale changes in codon bias, a codon-deoptimized A24R cDNA was chemically synthesized *de novo* (Biobasic). The synthetized A24R gene was inserted in a plasmid for the generation of rVV, including left and right (L and R, respectively) flanking A24R regions for homologous recombination in A24R locus and a poxviral early transcription terminator sequence (TTTTTGT) (pUCSP-A24cd). Then, the TagGFP2 gene cassette derived from pRB-TagGFP2 (69) was subcloned into pUCSP-A24cd using XhoI and BamHI restriction enzymes using standard molecular biology techniques to generate the plasmid pUCSP-A24cd-GFP, which was used to generate the recombinant v-A24Rcd. In addition, a plasmid containing the A24Rwt sequence was engineered (pUCSP-A24wt-GFP) to generate a revertant rVV expressing the wild-type version of A24R gene (v-A24Rre). To that end, A24Rwt was amplified by PCR and cloned into pUCSP-A24cd-GFP using SphI and XhoI restriction enzymes using standard molecular techniques. A24Rwt and A24Rcd were amplified by PCR using specific primers and cloned into a pCAGGS plasmid containing an HA epitope tag (70, 71) using SacI and SmaI restriction enzymes. All plasmids were generated using standard cloning techniques. Primers to generate the described plasmid constructs are available upon request. Plasmid constructs were verified by DNA sequencing (ACGT or Macrogen).

### Generation of rVV

Recombinant viruses were generated following infection/transfection approaches as previously described (19, 20, 69). Briefly, to generate the rVV expressing A24Rcd (v-A24Rcd), BSC-1 cells (6-well plate format, 10^6^ cells/well, triplicates) were infected at multiplicity of infection (MOI) of 0.05 plaque forming units (PFU)/cell with vaccinia WR (20, 69). After 1 h viral adsorption, virus was removed and cells were transfected with 2 µg of plasmid DNA pUCSP-A24Rcd using FuGeneHD (Promega) following manufacturer’s recommendations. After 48-72 h, cells were harvested, and recombinant v-A24Rcd was isolated from progeny virus by 5 rounds of plaque purification on BSC-1 cells, during which the plaques were screened for GFP expression. The small-plaque forming virus v-A24Rcd was plaque purified and amplified. v-A24Rcd and the plasmid pUCSP-A24wt-GFP were used to generate a revertant rVV expressing a wild-type A24R gene (v-A24Rre), following the same infection/transfection experimental protocol described above. Stable, large plaques were isolated in this case, and viral stocks were generated following amplification in BSC-1 cells and viral titers (PFU/ml) were determined by standard plaque assay.

### Transient cell transfections

To evaluate expression levels of A24Rwt and A24Rcd, human 293T cells (10^5^ cells, 12-well plates, triplicates) were transiently transfected using Lipofectamine 2000 (LPF2000, Invitrogen) with 1 µg of pCAGGS expression plasmids encoding C-terminal HA-tagged A24Rwt or A24Rcd. At 48 h post-transfection (p.t), protein expression levels were analyzed by immunofluorescence as indicated below. In addition, human 293T cells (10^6^ cells in 6-well plates, triplicates) were transiently co-transfected, using LPF2000, with pCAGGS expression plasmids versions of A24Rwt or A24Rcd proteins (2 µg) together with a pCAGGS plasmid encoding GFP (0.5 µg). At 48 h p.t., GFP expression was assessed under a fluorescence microscope, and cells were collected by centrifugation and lysed using RIPA buffer (72, 73). Protein expression levels were evaluated by Western blot as indicated below.

### SDS-PAGE electrophoresis and Western blot analysis

Total proteins from transfected or infected cell lysates were separated by denaturing electrophoresis in 10% SDS-polyacrylamide gels and transferred to nitrocellulose membranes. After transfer, membranes were blocked for 1 h with 10% dried skim milk in phosphate-buffered saline (PBS), 0.1% Tween 20 (T-PBS) and incubated overnight at 4°C with specific primary monoclonal (MAbs) or polyclonal (pAbs) antibodies: anti-HA tag (rabbit pAb; H6908, Sigma), anti-GFP (rabbit pAb; SC8334, Santa Cruz), anti-F13L (rat MAb 15B6, kindly provided by Dr. Gerhard Hiller), or anti-VV (in house-made rabbit pAb). Mouse MAb against actin (A1978, Sigma) was used as an internal loading control. Bound primary antibodies were detected with horseradish peroxidase (HRP)-conjugated secondary antibodies against the different species, and proteins were detected by chemiluminescence (Hyglo; Dennville Scientific), following the manufacturer’s recommendations. Western blots were quantified by densitometry using the software ImageJ (v1.46). Protein bands were normalized to the level of actin expression. The level of expression of the A24Rwt protein was considered 100% for comparison with the levels of expression of the A24Rcd construct.

### Immunofluorescence and fluorescence microcopy

To assess cellular GFP expression from transfected or infected cells using fluorescence, cells were directly imaged under a fluorescence microscope. To analyze A24Rwt or A24Rcd expression from transfected cells using indirect immunofluorescence, cells were fixed and permeabilized with 4% (vol/vol) formaldehyde and 0.5% (vol/vol) Triton X-100 (Sigma), respectively, before blocking with PBS containing 2.5% FBS for 1 h at room temperature. Thereafter, cells were incubated with an anti-HA epitope tag rabbit pAb (H6908, Sigma) and, after 3 consecutive washes with PBS, with a FITC-conjugated donkey anti-rabbit secondary antibody (Invitrogen). After 3 washes with PBS, cell nuclei were stained with DAPI (4′,6-diamindino-2-phenylindole). Representative images were obtained using a fluorescent microscope. To analyze A24Rwt or A24Rcd expression from infected cells by indirect immunofluorescence, infected cells on round coverslips in 24-well plates were washed twice with PBS and fixed by the addition of ice-cold 4% paraformaldehyde for 12 min. All subsequent incubations were carried out at room temperature. Cells were permeabilized by a 15 min incubation in PBS containing 0.1% Triton X-100 (Sigma). Cells were treated with PBS containing 0.1 M glycine for 5 min and incubated with primary antibodies diluted in PBS-20% FCS for 30 min followed by incubation with secondary antibodies diluted in PBS-20% FCS. DNA was stained with Hoechst by incubating cell on glass coverslips with 2 mg/ml bisbenzimide (Hoechst dye. Sigma) for 30 min. Antibodies used were rat MAb 19C2 (anti-B5, kindly provided by Dr. Gerhard Hiller), and anti-rat Alexa Fluor 488 (Invitrogen). Finally, cells were washed extensively with PBS, mounted with FluorSave reagent (Calbiochem), and observed by fluorescence microscopy.

### Plaque assays and crystal violet staining

To assess the plaque phenotype of the rVV, BSC-1 cells (6-well plate format, 10^6^ cells/well, triplicates) were infected with approximately 100-200 PFU per well. After 1 h infection, the medium was removed, and the infection was maintained for 48 h at 37°C under semisolid medium consisting of EMEM-2% FBS containing 0.7% methyl cellulose. Cell monolayers were subsequently fixed and stained using a solution of 5% (v/v) crystal violet in 20% methanol. After removal of the semisolid medium with crystal violet, plaques were photographed. The area of plaques was quantified using ImageJ.

### Virus growth kinetics

Growth kinetics were evaluated in BSC-1 cells (6-well plate format, 10^6^ cells/well, triplicates) infected at an MOI of 0.01. Viral replication at the indicated times post-infection (p.i.) (24, 48, 72, and 96 h), was evaluated by fluorescence microscopy. In addition, at each time point, cell culture supernatants were collected to determine viral titers using standard plaque assay followed by staining with a solution of 5% (v/v) crystal violet in 20% methanol. Microsoft Excel software and GraphPad Prism were used to determine the mean value and standard deviation (SD).

### Mouse experiments

All animal experiments were approved by the University of Rochester Institutional Biosafety (IBC) and Animal Care and Use (IACUC) Committees, which are in accordance with recommendations found in the Guide for the Care and Use of Laboratory Animals of the National Research Council (74). Six-to-eight-weeks-old female C57BL/6 mice were purchased from the National Cancer Institute (NCI) and maintained at the University of Rochester animal care facility under specific-pathogen-free conditions. Mice (n = 6/group) were anesthetized intraperitoneally (i.p.) with ketamine-xylazine (100 mg/kg ketamine and 10 mg/kg xylazine). For vaccination studies, mice were infected intranasally (i.n.) with the indicated PFU (10^4^, 10^5^, or 10^6^ PFU) and monitored each day for changes in morbidity (body weight loss) and mortality (survival) for 14 days. For challenge studies, twenty-two days after vaccination, mice were challenged with a lethal dose (10^4^ PFU) of parental VV WR and morbidity and survival were evaluated for 14 days. Mice losing more than 25% of their initial body weight were considered to reach their end point and were humanely euthanized with carbon dioxide (CO_2_) and confirmed by cervical dislocation.

### Genetic stability of rVV

To determine the genetic stability of v-A24Rcd, virus was serially passaged 10 times in BSC-1 cells (6-well plate format, 10^6^ cells/well) and incubated for 48 h until cytopathic effect (CPE) was observed. First passage was initiated using an MOI of 0.1. Inoculum of the successive passages were 1:20 dilutions of the cell extract from the previous passage. After 10 consecutive passages, P1 and P10 viruses were compared to WR and v-GFP by plaque assay and growth kinetics as indicated above.

## ACKNOWLEDGMENTS

We want to thank Dr. Gerhard Hiller for providing antibodies. This work was supported by grants E-700 RTA2014-00006, RTA2017-0066 from Ministerio de Economía y Competitividad and Ministerio de Ciencia, Innovación y Universidades as part of the Plan Estatal de Investigación Científica, Desarrollo e Innovación Tecnológica, and grant COV20-00901 703 from Instituto de Salud Carlos III (ISCIII). This research was funded by a “Ramon y Cajal” Incorporation grant (RYC-2017) from the Spanish Ministry of Science, Innovation and Universities to A.N. Research in LMS was partially funded by the Department of Defense (DoD) W81XWH1910496 and the National Institute of Health (NIH) R21 AI1135284 grants, and startup funding from the Department of Microbiology and Immunology at University of Rochester.

## Notes

### Competing Interest Statement

The authors have declared no competing interest.

