## Supplementary Figure 1 for "Vaccinia virus attenuation by codon deoptimization of the A24R gene for vaccine development"

**Supplementary Figure 1. Nucleotide sequence of wild type (top) and CD (bottom) VV A24R.**
Red indicates nucleotide changes introduced to CD VV A24R.

| | |
|---|---|
| **A24R** | ATG AAA AAA AAC ACT GAT TCA GAA ATG GAT CAA CGA CTA GGG TAT AAG TTT |
| **CD** | ATG AAA AAA AAT ACG GAT TCG GAA ATG GAT CAA CGT CTA GGT TAT AAA TTT |
| **A24R** | TTG GTG CCT GAT CCT AAA GCC GGA GTT TTT TAT AGA CCG TTA CAT TTC CAA |
| **CD** | CTA GTA CCG GAC CCG AAA GCG GGT GTA TTT TAT CGT CCG CTA CAT TTT CAA |
| **A24R** | TAT GTA TCG TAT TCT AAT TTT ATA TTG CAT CGA TTG CAT GAA ATC TTG ACC |
| **CD** | TAT GTA TCG TAT TCG AAT TTT ATA CTA CAT CGT CTA CAT GAA ATA CTA ACG |
| **A24R** | GTC AAG CGG CCA CTC TTA TCG TTT AAG AAT AAT ACA GAA CGA ATT ATG ATA |
| **CD** | GTA AAA CGT CCG CTA CTA TCG TTT AAA AAT AAT ACG GAA CGT ATA ATG ATA |
| **A24R** | GAA ATT AGC AAT GTT AAA GTG ACT CCT CCA GAT TAC TCA CCT ATA ATC GCG |
| **CD** | GAA ATA TCG AAT GTA AAA GTA ACG CCG CCG GAT TAT TCG CCG ATA ATA GCG |
| **A24R** | AGT ATT AAA GGT AAG AGT TAT GAT GCA TTA GCC ACG TTC ACT GTA AAT ATC |
| **CD** | TCG ATA AAA GGT AAA TCG TAT GAT GCG CTA GCG ACG TTT ACG GTA AAT ATA |
| **A24R** | TTT AAA GAG GTA ATG ACC AAA GAG GGT ATA TCC ATC ACT AAA ATA AGT AGT |
| **CD** | TTT AAA GAA GTA ATG ACG AAA GAA GGT ATA TCG ATA ACG AAA ATA TCG TCG |
| **A24R** | TAT GAG GGA AAA GAT TCT CAT TTG ATA AAA ATT CCG CTA CTA ATA GGA TAC |
| **CD** | TAT GAA GGT AAA GAT TCG CAT CTA ATA AAA ATA CCG CTA CTA ATA GGT TAT |
| **A24R** | GGG AAT AAA AAT CCA CTT GAT ACA GCC AAG TAT CTT GTT CCT AAT GTC ATA |
| **CD** | GGT AAT AAA AAT CCG CTA GAT ACG GCG AAA TAT CTA GTA CCG AAT GTA ATA |
| **A24R** | GGT GGA GTC TTT ATC AAT AAA CAA TCT GTC GAA AAA GTA GGA ATT AAT CTA |
| **CD** | GGT GGT GTA TTT ATA AAT AAA CAA TCG GTA GAA AAA GTA GGT ATA AAT CTA |
| **A24R** | GTA GAA AAG ATT ACA ACA TGG CCA AAA TTT AGG GTT GTT AAG CCA AAC TCA |
| **CD** | GTA GAA AAA ATA ACG ACG TGG CCG AAA TTT CGT GTA GTA AAA CCG AAT TCG |
| **A24R** | TTC ACT TTC TCG TTT TCC TCC GTA TCC CCT CCT AAT GTA TTA CCG ACA AGA |
| **CD** | TTT ACG TTT TCG TTT TCG TCG GTA TCG CCG CCG AAT GTA CTA CCG ACG CGT |
| **A24R** | TAT CGC CAT TAC AAG ATA TCT CTG GAT ATA TCA CAA TTG GAA GCG TTG AAT |
| **CD** | TAT CGT CAT TAT AAA ATA TCG CTA GAT ATA TCG CAA CTA GAA GCG CTA AAT |
| **A24R** | ATA TCA TCG ACA AAG ACA TTT ATA ACG GTC AAT ATT GTT TTG CTG TCT CAA |
| **CD** | ATA TCG TCG ACG AAA ACG TTT ATA ACG GTA AAT ATA GTA CTA CTA TCG CAA |
| **A24R** | TAT TTA TCT AGA GTG AGT CTA GAA TTC ATT AGA CGT AGT TTA TCA TAC GAT |
| **CD** | TAT CTA TCG CGT GTA TCG CTA GAA TTT ATA CGT CGT TCG CTA TCG TAT GAT |
| **A24R** | ATG CCT CCA GAA GTT GTC TAT CTA GTA AAC GCG ATA ATA GAT AGT GCT AAA |
| **CD** | ATG CCG CCG GAA GTA GTA TAT CTA GTA AAT GCG ATA ATA GAT TCG GCG AAA |

| | |
|---|---|
| **A24R** | CGA ATT ACT GAA TCT ATT ACT GAC TTT AAT ATT GAT ACA TAC ATT AAT GAC |
| **CD** | CGT ATA ACG GAA TCG ATA ACG GAT TTT AAT ATA GAT ACG TAT ATA AAT GAT |
| **A24R** | CTG GTG GAA GCT GAA CAC ATT AAA CAA AAA TCT CAG TTA ACG ATC AAC GAG |
| **CD** | CTA GTA GAA GCG GAA CAT ATA AAA CAA AAA TCG CAA CTA ACG ATA AAT GAA |
| **A24R** | TTC AAA TAT GAA ATG CTG CAT AAC TTT TTA CCT CAT ATG AAC TAT ACA CCC |
| **CD** | TTT AAA TAT GAA ATG CTA CAT AAT TTT CTA CCG CAT ATG AAT TAT ACG CCG |
| **A24R** | GAT CAA CTA AAG GGA TTT TAT ATG ATA TCT TTA CTA AGA AAG TTT CTC TAC |
| **CD** | GAT CAA CTA AAA GGT TTT TAT ATG ATA TCG CTA CTA CGT AAA TTT CTA TAT |
| **A24R** | TGT ATC TAC CAC ACT TCT AGA TAT CCA GAT AGA GAT TCG ATG GTT TGT CAT |
| **CD** | TGT ATA TAT CAT ACG TCG CGT TAT CCG GAT CGT GAT TCG ATG GTA TGT CAT |
| **A24R** | CGC ATC CTA ACG TAC GGC AAA TAT TTT GAG ACG TTG GCA CAT GAT GAA TTA |
| **CD** | CGT ATA CTA ACG TAT GGT AAA TAT TTT GAA ACG CTA GCG CAT GAT GAA CTA |
| **A24R** | GAG AAT TAC ATA GGC AAC ATC CGA AAC GAT ATC ATG AAC AAT CAC AAG AAC |
| **CD** | GAA AAT TAT ATA GGT AAT ATA CGT AAT GAT ATA ATG AAT AAT CAT AAA AAT |
| **A24R** | AGA GGC ACT TAC GCG GTA AAC ATT CAT GTA CTA ACA ACT CCC GGA CTT AAT |
| **CD** | CGT GGT ACG TAT GCG GTA AAT ATA CAT GTA CTA ACG ACG CCG GGT CTA AAT |
| **A24R** | CAC GCG TTT TCT AGC TTA TTG AGT GGA AAG TTC AAA AAG TCA GAC GGT AGT |
| **CD** | CAT GCG TTT TCG TCG CTA CTA TCG GGT AAA TTT AAA AAA TCG GAT GGT TCG |
| **A24R** | TAT CGA ACA CAT CCT CAC TAT TCA TGG ATG CAG AAT ATT TCT ATT CCT AGG |
| **CD** | TAT CGT ACG CAT CCG CAT TAT TCG TGG ATG CAA AAT ATA TCG ATA CCG CGT |
| **A24R** | AGT GTT GGA TTT TAT CCG GAT CAA GTA AAG ATT TCA AAG ATG TTT TCT GTC |
| **CD** | TCG GTA GGT TTT TAT CCG GAT CAA GTA AAA ATA TCG AAA ATG TTT TCG GTA |
| **A24R** | AGA AAA TAC CAT CCA AGT CAA TAT CTT TAC TTT TGT TCA TCG GAC GTT CCG |
| **CD** | CGT AAA TAT CAT CCG TCG CAA TAT CTA TAT TTT TGT TCG TCG GAT GTA CCG |
| **A24R** | GAA AGA GGT CCT CAG GTA GGT TTA GTA TCT CAA TTG TCT GTC TTG AGT TCC |
| **CD** | GAA CGT GGT CCG CAA GTA GGT CTA GTA TCG CAA CTA TCG GTA CTA TCG TCG |
| **A24R** | ATT ACA AAT ATA CTA ACG TCT GAG TAT TTG GAT TTG GAA AAG AAA ATT TGT |
| **CD** | ATA ACG AAT ATA CTA ACG TCG GAA TAT CTA GAT CTA GAA AAA AAA ATA TGT |
| **A24R** | GAG TAT ATC AGA TCA TAT TAT AAA GAT GAT ATA AGT TAC TTT GAA ACA GGA |
| **CD** | GAA TAT ATA CGT TCG TAT TAT AAA GAT GAT ATA TCG TAT TTT GAA ACG GGT |
| **A24R** | TTT CCA ATC ACT ATA GAA AAT GCT CTA GTC GCA TCT CTT AAT CCA AAT ATG |
| **CD** | TTT CCG ATA ACG ATA GAA AAT GCG CTA GTA GCG TCG CTA AAT CCG AAT ATG |
| **A24R** | ATA TGT GAT TTT GTA ACT GAC TTT AGA CGT AGA AAA CGG ATG GGA TTC TTC |
| **CD** | ATA TGT GAT TTT GTA ACG GAT TTT CGT CGT CGT AAA CGT ATG GGT TTT TTT |

| | |
|---|---|
| **A24R** | GGT AAC TTG GAG GTA GGT ATT ACT TTA GTT AGG GAT CAC ATG AAT GAA ATT |
| **CD** | GGT AAT CTA GAA GTA GGT ATA ACG CTA GTA CGT GAT CAT ATG AAT GAA ATA |
| **A24R** | CGC ATT AAT ATT GGA GCG GGA AGA TTA GTC AGA CCA TTC TTG GTT GTG GAT |
| **CD** | CGT ATA AAT ATA GGT GCG GGT CGT CTA GTA CGT CCG TTT CTA GTA GTA GAT |
| **A24R** | AAC GGA GAG CTC ATG ATG GAT GTG TGT CCG GAG TTA GAA AGC AGA TTA GAC |
| **CD** | AAT GGT GAA CTA ATG ATG GAT GTA TGT CCG GAA CTA GAA TCG CGT CTA GAT |
| **A24R** | GAC ATG ACA TTC TCT GAC ATT CAG AAA GAG TTT CCG CAT GTC ATC GAA ATG |
| **CD** | GAT ATG ACG TTT TCG GAT ATA CAA AAA GAA TTT CCG CAT GTA ATA GAA ATG |
| **A24R** | GTA GAT ATA GAA CAA TTT ACT TTT AGT AAC GTA TGT GAA TCG GTT CAA AAA |
| **CD** | GTA GAT ATA GAA CAA TTT ACG TTT TCG AAT GTA TGT GAA TCG GTA CAA AAA |
| **A24R** | TTT AGA ATG ATG TCA AAG GAT GAA AGA AAG CAA TAC GAT TTA TGT GAC TTT |
| **CD** | TTT CGT ATG ATG TCG AAA GAT GAA CGT AAA CAA TAT GAT CTA TGT GAT TTT |
| **A24R** | CCT GCC GAA TTT AGA GAT GGA TAT GTG GCA TCT TCA TTA GTG GGA ATC AAT |
| **CD** | CCG GCG GAA TTT CGT GAT GGT TAT GTA GCG TCG TCG CTA GTA GGT ATA AAT |
| **A24R** | CAC AAT TCT GGA CCC AGA GCT ATT CTT GGA TGT GCT CAA GCT AAA CAA GCT |
| **CD** | CAT AAT TCG GGT CCG CGT GCG ATA CTA GGT TGT GCG CAA GCG AAA CAA GCG |
| **A24R** | ATC TCT TGT CTG AGT TCG GAT ATA CGA AAT AAA ATA GAC AAT GGA ATT CAT |
| **CD** | ATA TCG TGT CTA TCG TCG GAT ATA CGT AAT AAA ATA GAT AAT GGT ATA CAT |
| **A24R** | TTG ATG TAT CCA GAG AGG CCA ATC GTG ATT AGT AAG GCT TTA GAA ACT TCA |
| **CD** | CTA ATG TAT CCG GAA CGT CCG ATA GTA ATA TCG AAA GCG CTA GAA ACG TCG |
| **A24R** | AAG ATT GCG GCT AAT TGC TTC GGC CAA CAT GTT ACT ATA GCA TTA ATG TCG |
| **CD** | AAA ATA GCG GCG AAT TGT TTT GGT CAA CAT GTA ACG ATA GCG CTA ATG TCG |
| **A24R** | TAC AAA GGT ATC AAT CAA GAG GAT GGA ATT ATC ATC AAA AAA CAA TTT ATT |
| **CD** | TAT AAA GGT ATA AAT CAA GAA GAT GGT ATA ATA ATA AAA AAA CAA TTT ATA |
| **A24R** | CAG AGA GGC GGT CTC GAT ATA GTT ACC GCA AAG AAA CAT CAA GTA GAA ATT |
| **CD** | CAA CGT GGT GGT CTA GAT ATA GTA ACG GCG AAA AAA CAT CAA GTA GAA ATA |
| **A24R** | CCG TTG GAA AAC TTT AAT AAC AAA GAA AGA GAT AGG TCT AAC GCC TAT TCA |
| **CD** | CCG CTA GAA AAT TTT AAT AAT AAA GAA CGT GAT CGT TCG AAT GCG TAT TCG |
| **A24R** | AAA TTA GAA AGT AAT GGA TTA GTT AGA CTG AAT GCT TTC TTG GAA TCC GGA |
| **CD** | AAA CTA GAA TCG AAT GGT CTA GTA CGT CTA AAT GCG TTT CTA GAA TCG GGT |
| **A24R** | GAC GCT ATG GCA CGA AAT ATC TCA TCA AGA ACT CTT GAA GAT GAT TTT GCT |
| **CD** | GAT GCG ATG GCG CGT AAT ATA TCG TCG CGT ACG CTA GAA GAT GAT TTT GCG |
| **A24R** | AGA GAT AAT CAG ATT AGC TTC GAT GTT TCC GAG AAA TAT ACC GAT ATG TAC |
| **CD** | CGT GAT AAT CAA ATA TCG TTT GAT GTA TCG GAA AAA TAT ACG GAT ATG TAT |

```
A24R    AAA TCT CGC GTT GAA CGA GTA CAA GTA GAA CTT ACT GAC AAA GTT AAG GTA
CD      AAA TCG CGT GTA GAA CGT GTA CAA GTA GAA CTA ACG GAT AAA GTA AAA GTA

A24R    CGA GTA TTA ACC ATG AAA GAA AGA AGA CCC ATT CTA GGA GAT AAA TTT ACC
CD      CGT GTA CTA ACG ATG AAA GAA CGT CGT CCG ATA CTA GGT GAT AAA TTT ACG

A24R    ACT AGA ACG AGT CAA AAG GGA ACA GTC GCG TAT GTC GCG GAT GAA ACG GAA
CD      ACG CGT ACG TCG CAA AAA GGT ACG GTA GCG TAT GTA GCG GAT GAA ACG GAA

A24R    CTT CCA TAC GAC GAA AAT GGT ATC ACA CCA GAT GTC ATT ATT AAT TCT ACA
CD      CTA CCG TAT GAT GAA AAT GGT ATA ACG CCG GAT GTA ATA ATA AAT TCG ACG

A24R    TCC ATC TTC TCT AGA AAA ACT ATA TCT ATG TTG ATA GAG GTT ATT TTA ACA
CD      TCG ATA TTT TCG CGT AAA ACG ATA TCG ATG CTA ATA GAA GTA ATA CTA ACG

A24R    GCC GCA TAT TCT GCT AAG CCG TAC AAC AAT AAG GGA GAA AAC CGA CCT GTC
CD      GCG GCG TAT TCG GCG AAA CCG TAT AAT AAT AAA GGT GAA AAT CGT CCG GTA

A24R    TGT TTT CCT AGT AGT AAC GAA ACA TCC ATC GAT ACA TAT ATG CAA TTC GCT
CD      TGT TTT CCG TCG TCG AAT GAA ACG TCG ATA GAT ACG TAT ATG CAA TTT GCG

A24R    AAA CAA TGT TAT GAG CAT TCA AAT CCG AAA TTG TCC GAT GAA GAA TTA TCG
CD      AAA CAA TGT TAT GAA CAT TCG AAT CCG AAA CTA TCG GAT GAA GAA CTA TCG

A24R    GAT AAA ATC TTT TGT GAA AAG ATT CTC TAT GAT CCT GAA ACG GAT AAG CCT
CD      GAT AAA ATA TTT TGT GAA AAA ATA CTA TAT GAT CCG GAA ACG GAT AAA CCG

A24R    TAT GCA TCC AAA GTA TTT TTT GGA CCA ATT TAT TAC TTG CGT CTG AGG CAT
CD      TAT GCG TCG AAA GTA TTT TTT GGT CCG ATA TAT TAT CTA CGT CTA CGT CAT

A24R    TTA ACT CAG GAC AAG GCA ACC GTT AGA TGT AGA GGT AAA AAG ACG AAG CTC
CD      CTA ACG CAA GAT AAA GCG ACG GTA CGT TGT CGT GGT AAA AAA ACG AAA CTA

A24R    ATT AGA CAG GCG AAT GAG GGA CGA AAA CGT GGA GGA GGT ATC AAG TTC GGA
CD      ATA CGT CAA GCG AAT GAA GGT CGT AAA CGT GGT GGT GGT ATA AAA TTT GGT

A24R    GAA ATG GAG AGA GAC TGT TTA ATA GCG CAT GGC GCA GCC AAT ACT ATT ACA
CD      GAA ATG GAA CGT GAT TGT CTA ATA GCG CAT GGT GCG GCG AAT ACG ATA ACG

A24R    GAA GTT TTA AAA GAC TCA GAA GAG GAT TAT CAA GAT GTG TAT GTT TGT GAA
CD      GAA GTA CTA AAA GAT TCG GAA GAA GAT TAT CAA GAT GTA TAT GTA TGT GAA

A24R    AAT TGT GGA GAC ATA GCA GCA CAA ATC AAG GGT ATT AAT ACA TGT CTT AGA
CD      AAT TGT GGT GAT ATA GCG GCG CAA ATA AAA GGT ATA AAT ACG TGT CTA CGT

A24R    TGT TCA AAA CTT AAT CTC TCT CCT CTC TTA ACA AAA ATT GAT ACC ACG CAC
CD      TGT TCG AAA CTA AAT CTA TCG CCG CTA CTA ACG AAA ATA GAT ACG ACG CAT


A24R    GTA TCT AAA GTA TTT CTT ACT CAA ATG AAC GCC AGA GGC GTA AAA GTC AAA
CD      GTA TCG AAA GTA TTT CTA ACG CAA ATG AAT GCG CGT GGT GTA AAA GTA AAA
```

| | |
|---|---|
| **A24R** | TTA GAT TTC GAA CGA AGG CCT CCT TCG TTT TAT AAA CCA TTA GAT AAA GTT |
| **CD** | CTA GAT TTT GAA CGT CGT CCG CCG TCG TTT TAT AAA CCG CTA GAT AAA GTA |

| | |
|---|---|
| **A24R** | GAT CTC AAG CCG TCT TTT CTG GTG TAA |
| **CD** | GAT CTA AAA CCG TCG TTT CTA GTA TAA |
